# Computational genomic discovery of diverse gene clusters harboring Fe-S flavoenzymes in anaerobic gut microbiota

**DOI:** 10.1101/2020.03.06.976126

**Authors:** Victòria Pascal Andreu, Michael A. Fischbach, Marnix H. Medema

## Abstract

The gut contains an enormous diversity of simple as well as complex molecules from highly diverse food sources as well as host-secreted molecules. This presents a large metabolic opportunity for the gut microbiota, but little is known on how gut microbes are able to catabolize this large chemical diversity. Recently, Fe-S flavoenzymes were found to be key in the transformation of bile acids, catalysing the key step in the 7*α*-dehydroxylation pathway that allows gut bacteria to transform cholic acid (CA) into deoxycholic acid (DCA), an exclusively microbe-derived molecule with major implications for human health. While this enzyme family has also been implicated in a limited number of other catalytic transformations, little is known about the extent to which it is of more global importance in gut microbial metabolism. Here, we use large-scale computational genomic analysis to show that this enzyme superfamily has undergone a remarkable expansion in Clostridiales, and occurs throughout a diverse array of >1,000 different families of putative metabolic gene clusters. Analysis of the enzyme content of these gene clusters suggests that they encode pathways with a wide range of predicted substrate classes, including saccharides, amino acids/peptides and lipids. Altogether, these results indicate a potentially important role of this protein superfamily in the human gut, and our dataset provides significant opportunities for the discovery of novel pathways that may have significant effects on human health.

## Introduction

The gene set of the human gut microbiota vastly exceeds the human gene repertoire^1,2^, which allows microbes to complement human metabolism by degrading undigested polysaccharides, lipids, and peptides that reach the large intestine^3^. For instance, saccharolytic bacteria can ferment carbohydrates to produce short chain fatty acids (SCFAs), beneficial metabolites that promote health^4^. Nevertheless, gut bacteria also produce many molecules involved in microbe-microbe interactions and microbe-host interactions that can have detrimental effects instead^5^. An example of a harmful diet-derived metabolite is trimethylamine (TMA), an amine that can be synthesize from choline or carnitine by certain gut bacteria, and which has been associated with cardiovascular and renal disease^6^. Thus, the identification of these molecules and the elucidation of their producing pathways is crucial to assess the causes and consequences of certain microbiome-associated phenotypes.

Another example of exclusively microbiome-derived molecules are the secondary bile acids (BA): deoxycholic acid (DCA) and lithocholic acid (LCA). While the primary bile acids cholic acid (CA) and chenodeoxycholic acid (CDCA) are synthesized by the liver^7^, they are transformed into DCA or LCA by colonic bacteria during enterohepatic circulation^8^. These molecules have been proposed to act as inhibitors of *C. difficile* outgrowth^9^, as well as to induce the development of colon cancer^10,11^ and cholesterol gallstone disease^12^. The main bacterial pathway in charge of this reaction occurs through 7*α*-dehydroxylation, a multi-step biochemical reaction that can be accomplished by bacteria harboring the bile-acid-inducible (*bai*) operon^13^. Most of the bacteria capable of carrying out this reaction are anaerobes that are part of the *Clostridium* cluster XIVa^13^. The pathway encoded by the *bai* operon was recently elucidated by Funabashi & Grove *et. al*^14^. The authors showed that the key step is performed by the BaiCD enzyme, an Fe-S flavoenzyme that oxidizes 3-oxo-cholyl-CoA to 3-oxo-4,5-dehydrocholyl-CoA. Later, BaiH (also an Fe-S flavoenzyme), BaiCD and BaiA2 act again on the molecule to finally produce DCA. Importantly, the participation of Fe-S flavoenzymes in the key reductive steps of the pathway is consistent with a role for this pathway in using primary bile acids as terminal electron acceptors for an anaerobic electron transport pathway, constituting a unique metabolic niche within the gut community.

In addition to the BaiCD and BaiH enzymes, a few other members of the Fe-S flavoenzyme superfamily have been previously shown to play similar crucial roles in the redox metabolism of catalytic transformations. For instance, L-phenylalanine fermentation via a Stickland reaction, where cinnamate is reduced to 3-phenylpropionate, has been shown to be performed by a cinnamate reductase that is a member of the Fe-S flavoenzyme superfamily^15^. Additional Fe-S flavoenzyme representatives with experimentally characterized functions include a 2,4-dienoyl-CoA reductase implicated in fatty acid beta-oxidation^16^ as well as a trimethylamine dehydrogenase involved in trimethylamine degradation^17^.

The fact that these enzymes have been shown to facilitate the shuttling of electrons from the membrane to diverse organic terminal electron acceptors, enabling an anaerobic electron transport chain, led us to hypothesize that they might play a more widespread role in the microbial catabolism of the diversity of complex substrates available in the gut. Here, we provide an in-depth computational genomic study of the Fe-S flavoenzyme superfamily and its presence across genomes of gut microbiome-related bacteria. Using phylogenomic analyses, we show that this enzyme superfamily comprises a large sequence diversity and has particularly undergone strong evolutionary expansion in the class Clostridia, emphasizing its possible implication on different metabolic reactions in the gut. Large amounts of strain-level variation between genomes indicates that the pathways involved likely facilitate ecologically specialized functions. Finally, analysis of the enzyme content in their surrounding operons and gene clusters uncovers a wide array of putative catabolic gene clusters associated with the breakdown of diverse substrates, which provides a rich resource for uncovering novel pathways in the human microbiome and beyond.

## Results & Discussion

### The Fe-S flavoenzymes phylogeny includes many unexplored clades involved in many catalytic reactions

In order to assess the distribution of the Fe-S flavoenzymes along the bacterial kingdom, we scanned 111,651 bacterial genomes (see Methods: *Identification of members of the Fe-S flavoenzyme superfamily*). Across this dataset, we identified 49,870 Fe-S flavoenzymes that belong to more than twenty bacterial phyla. Some bacterial genomes encode remarkable numbers of Fe-S flavoenzymes: e.g., the genome of the Firmicute *Sporobacter thermitidis* encodes no fewer than 15 Fe-S flavoenzymes. There are also representatives from Actinobacteria (including *Eggerthella sp. YY7918* and *Rhodococcus sp. SC4*) that possess 9 different flavoenzymes. On average, Firmicutes have 1.81 ± 1.18 protein copies per genome, whereas Proteobacteria and Actinobacteria have 1.51 ± 0.83 and 1.10 ± 0.50 copies respectively. These numbers exclude genomes that do not encode for any flavoenzyme, representing 89%, 55.5% and 46.8% of the genomes from Firmicutes, Proteobacteria and Actinobacteria respectively. Together, these data suggest that Fe-S flavoenzymes may play important roles in multiple bacterial phyla, and particularly in Firmicutes.

To contextualize these quantities of Fe-S flavoenzyme genes with their evolutionary history and functional diversification, we then conducted a comprehensive phylogenetic analysis of this enzyme family. We obtained a collection of 8,097 non-redundant Fe-S flavoenzymes by selecting representatives from MMseqs2^18^-derived sequence clusters. From this dataset, we constructed an approximate maximum likelihood phylogeny (Figure 1; see also Methods: *Construction of an Fe-S flavoenzyme protein similarity network*). Within this phylogeny, we were able to annotate eight clades based on experimentally characterized proteins documented in UniProt to date; the vast majority of clades have unknown functions. Given the size of this protein superfamily, the number of unexplored clades and its functional diversity, this phylogeny constitutes a road map to uncover novel clades of enzymes that could potentially aid in the characterization of yet undiscovered catabolic pathways.

**FIG 1.**
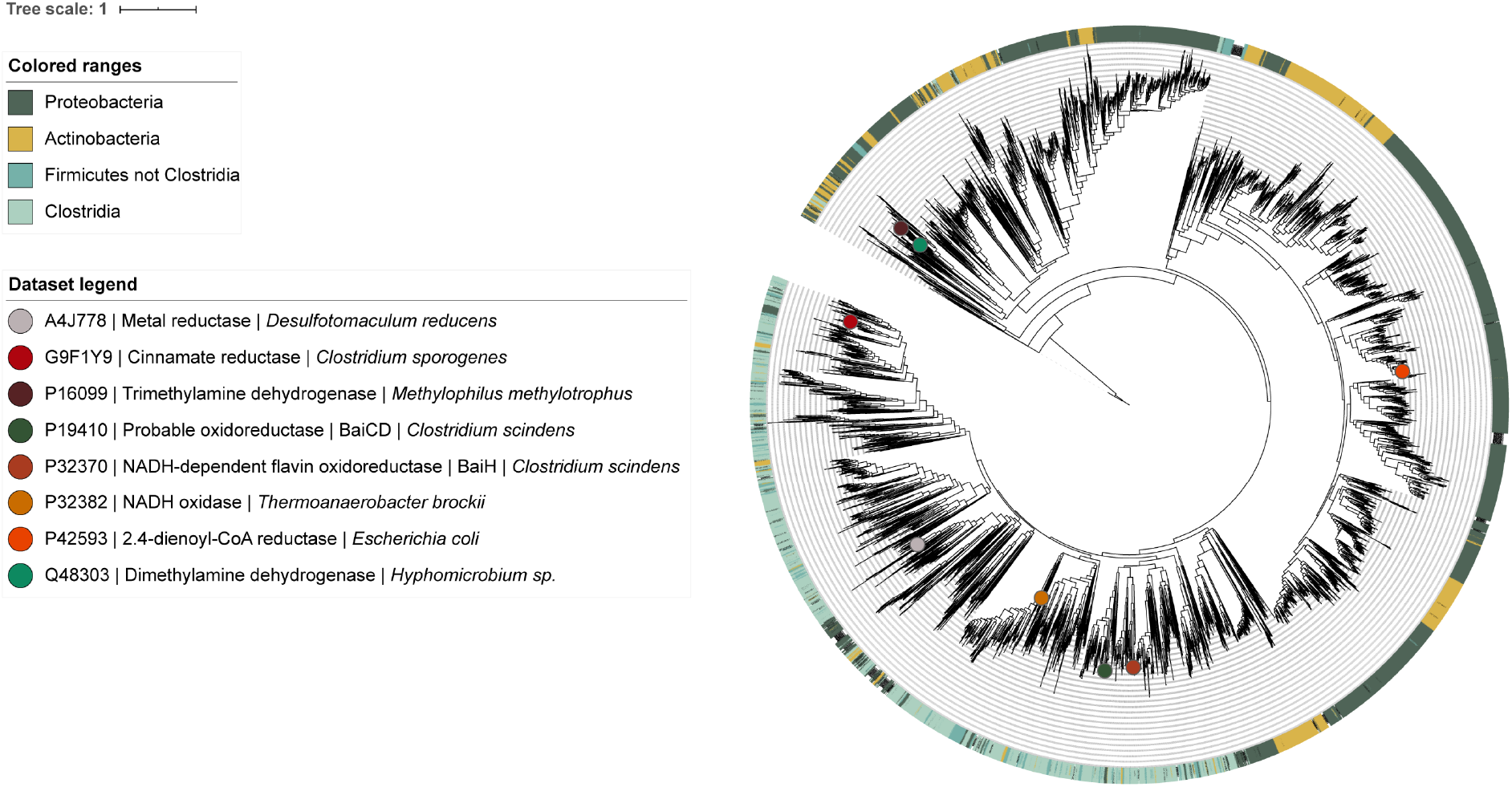
Fe-S Flavoenzyme superfamily phylogeny. The phylogeny is constructed from 8,105 non-redundant protein sequences across the bacterial kingdom, including the eight experimentally characterized proteins from Uniprot (https://www.uniprot.org/uniprot/?query=PF07992+and+PF00724+and+reviewed%3Ayes&sort=score, accessed at 01/02/2020). All sequences used as input harbor both the Oxidored_FMN and Pyr_redox_2 Pfam domains. The eight experimentally characterized proteins from Uniprot are indicated as circles on the tree to functionally annotate clades.

### An evolutionary expansion of Fe-S flavoenzyme in Clostridia

As reported in the previous section, the phylum Firmicutes stands out for the high Fe-S flavoenzyme copy numbers found in genomes of the species that belong to it. For this reason, we investigated the flavoenzymes copy number variation in *Clostridium* along with other microbial taxa commonly found in the gut: Firmicutes, Bacteroidetes, Actinobacteria and Proteobacteria. The phylogenetic analysis already suggested a large evolutionary expansion of Fe-S flavoenzymes in Clostridia, with this bacterial class occupying about a third of the non-redundant superfamily phylogeny. Indeed, quantitative analysis of the Clostridia class confirms that it has undergone a massive expansion when comparing the Fe-S flavoenzyme copy number with other taxa (Figure 2). The Actinobacteria and Bacteroidetes phyla seem to harbour considerably fewer Fe-S flavoenzymes. As expected, we found that among genomes from Clostridia, there is a high percentage that encode at least one Fe-S flavoenzyme (71.58%). In contrast, less than half of the genomes in our database belonging to Enterobacteriaceae, Bacilli and *Bacteroides* code for a flavoenzyme (49.92%, 7.64% and 3.19% respectively). Strikingly, *Eggerthell*a seems to be an exception to this pattern, as 10 genomes from different *Eggerthella* species encode 38 Fe-S flavoenzymes, with 71.43% of the genomes with at least one count.

**FIG 2.**
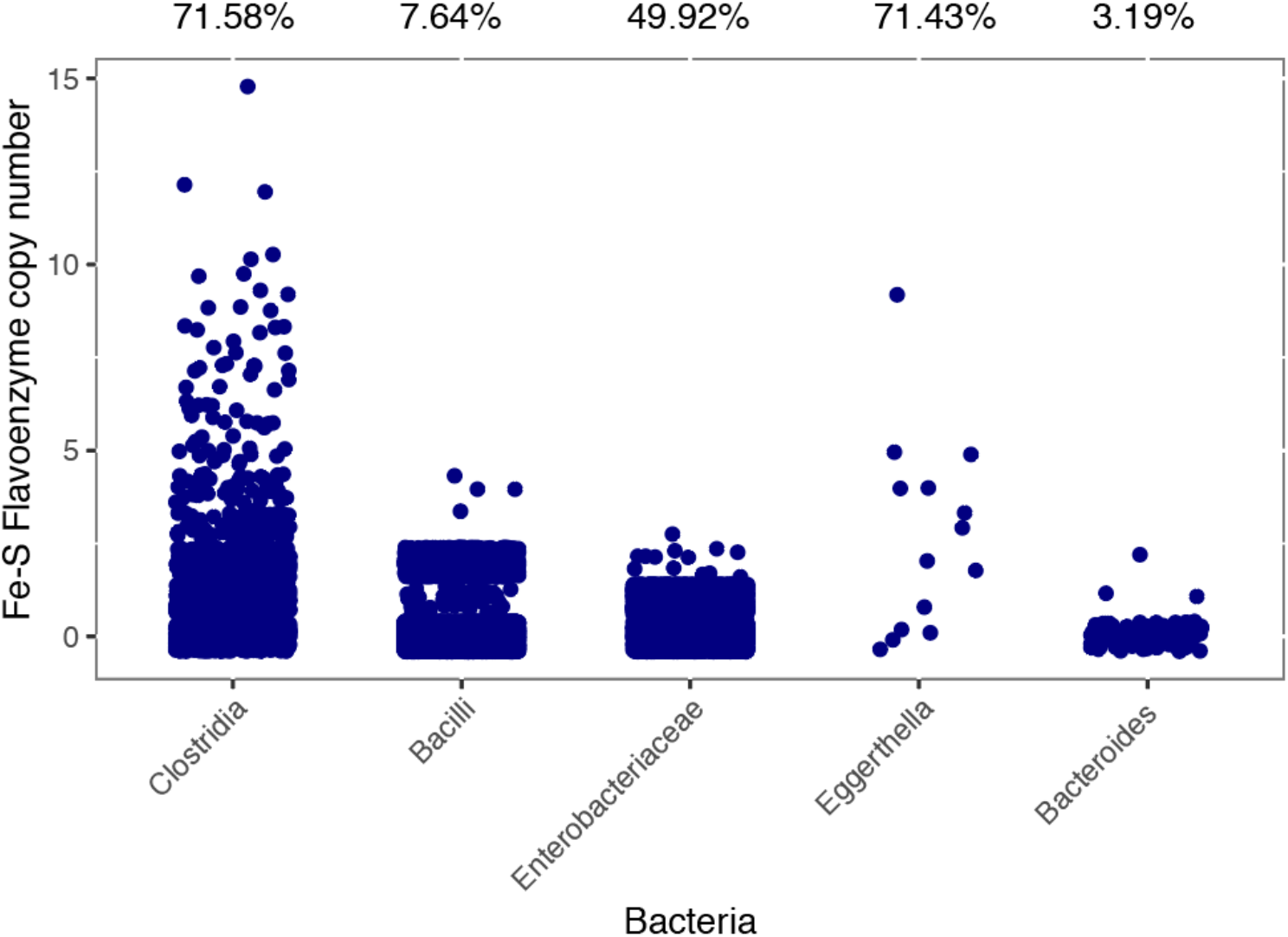
Expansion of the Fe-S flavoenzyme superfamily in Clostridia when compared to other gut-related taxa. Scatterplot representing the Fe-S flavoenzyme copy number across some gut-related bacteria. The taxonomic groups cover the five major phyla from the gut: Firmicutes (Clostridia and Bacilli), Proteobacteria (Enterobacteriaceae), Actinobacteria (*Eggerthella*) and Bacteroidetes (*Bacteroides*). The taxa picked are the ones found to possess at least one copy of flavoenzyme-encoding genes. The percentages on top represent how many of the genomes harbour at least one Fe-S flavoenzyme coding gene out of the total number of genomes from that taxon.

### Unexpected amounts of strain variation of flavoenzyme counts between genomes

In microbial ecosystems, strain-specific traits often confer different characteristics to otherwise almost identical bacteria, which allow them to thrive in specific conditions^19,20,21^. Therefore, we assessed Fe-S flavoenzyme copy number variation in order to know whether such enzymes are responsible for specialized functionalities that allow microbes to adapt to unique ecological niches. Indeed, we found that Fe-S flavoenzyme copy number distribution along Clostridia genomes often shows signs of strain-specificity, as can be seen in *Clostridioides difficile* clade and *Clostridia* clade II (Figure 3). In the Clostridia phylogeny (Figure 3), the number of copies ranges from 1 to 15 per genome, with a *Sporobacter thermitidis* genome being the one with highest counts. Another bacterium with a high Fe-S flavoenzyme copy number is *C. scindens* (Clostridia clade I), whose genome encodes up to 10 distinct members of the family (including BaiCD and BaiH). In comparison, other gut-related genera such as *Faecalibacterium* (clade IV, ~12 o’clock), *Roseburia* (clade I, ~5 o’clock) and *Anaerostipes* (clade I, ~4 o’clock) show quite different Fe-S flavoenzyme copy number profiles, harboring on average 0.3, 0.6 and 2.4 copies per genome, respectively. Overall, these results confirm that indeed, for specific clades of Clostridia, Fe-S flavoenzymes are likely to play an important role in their primary metabolism. Moreover, they evince the importance of profiling the microbiome at high (or strain) resolution; 16S amplicon sequencing (and especially OTU-level clustering) would not be able to distinguish between bacteria with different metabolic repertoires. The differential metabolic capabilities of various strains are often due to very subtle changes in their genomes^22^. Therefore, studying these bacterial strains that differ in terms of their Fe-S flavoenzyme copy numbers, as well as the genomic contexts where these enzymes are encoded, could help to uncover strain-specific traits that play a role in conferring microbiome-associated host phenotypes.

**FIG 3.**
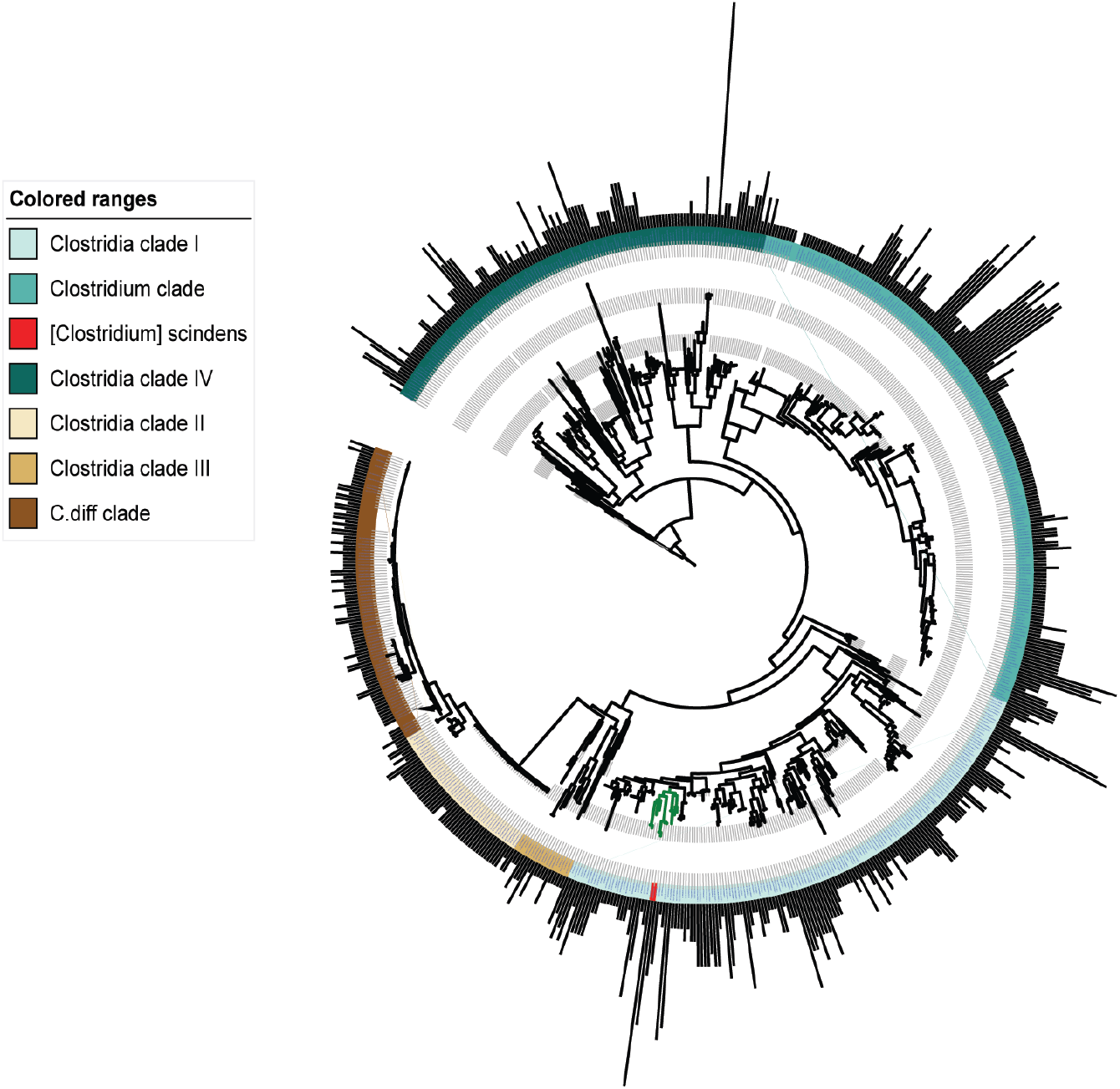
Strain- and species-specific copy numbers of Fe-S flavoenzymes across Clostridia. A circular phylogenetic tree showing bacterial strains in the class Clostridia. The outer ring of bars represents the copy number of Fe-S flavoenzymes in each strain, indicating a high degree of variability among strains and species. The phylogeny includes closely related strains whose genomes encode diverse numbers of Fe-S flavoenzymes, ranging from 1 to 15. Colored strips represent the taxonomic entities found within the class Clostridia. Two clades have been collapsed in order to remove almost identical genomes in the tree (shown as a discontinuity in the bar plots): the Clostridium difficile clade, Clostridia clade II and Clostridium clade.

### Analysis of gene neighborhoods leads to the identification of a wide range of families of putative catabolic gene clusters involved in breakdown of diverse biomolecules

We next explored the likely functionalities of flavoenzymes in Clostridia, by investigating the genomic contexts of the flavoenzyme-coding genes. With that aim, we gathered a collection of 3,158 non-redundant gene clusters, all of them having in common the presence of a Fe-S flavoenzyme-coding gene (see Methods; *Analysis of the genomic context of Fe-S flavoenzymes*). These gene clusters could be grouped into 1,052 Gene Cluster Families (GCFs). Interestingly, Fe-S flavoenzymes are found in a strikingly diverse array of GCFs. The examination of the corresponding gene cluster-coding proteins helped identifying more than 30 relevant Pfam domains that we categorized to be involved in 4 major types of metabolism (see **Table S1**). Thus, the profiling of this collection based on the presence of these protein domains within Fe-S-flavoenzyme-encoding gene clusters allowed to predict putative functions for 200 GCFs. Specifically, 80 distinct GCFs are predicted to function in saccharide catabolism, 83 in peptide/amino acid catabolism, 28 in nucleotide catabolism and 14 in lipid catabolism. In Figure 4, we highlight ten gene clusters plus the *bai* operon to show examples of distinct cluster architectures that, based on their enzyme-coding gene contents, we predict to be capable of processing a diverse array of substrates. Importantly, most of the Fe-S-flavoenzyme-containing gene clusters also harbor a major facilitator superfamily transporter, indicating that they might process a diffusible substrate; also, many include proteins with flavodoxin or electron transfer flavoprotein domains, consistent with the possibility that these pathways are coupled to electron transport chains. Overall, these findings suggest that Fe-S flavoenzymes play a far more expansive role in anaerobic metabolism in the human gut than was previously known. Moreover, the diversity in function manifests that these oxidoreductases may be in charge of catalyzing the transformation of a huge variety of substrates, conferring to these bacteria specialized primary metabolic capabilities that may allow them to colonize specific micro-niches in the gut.

**FIG 4.**
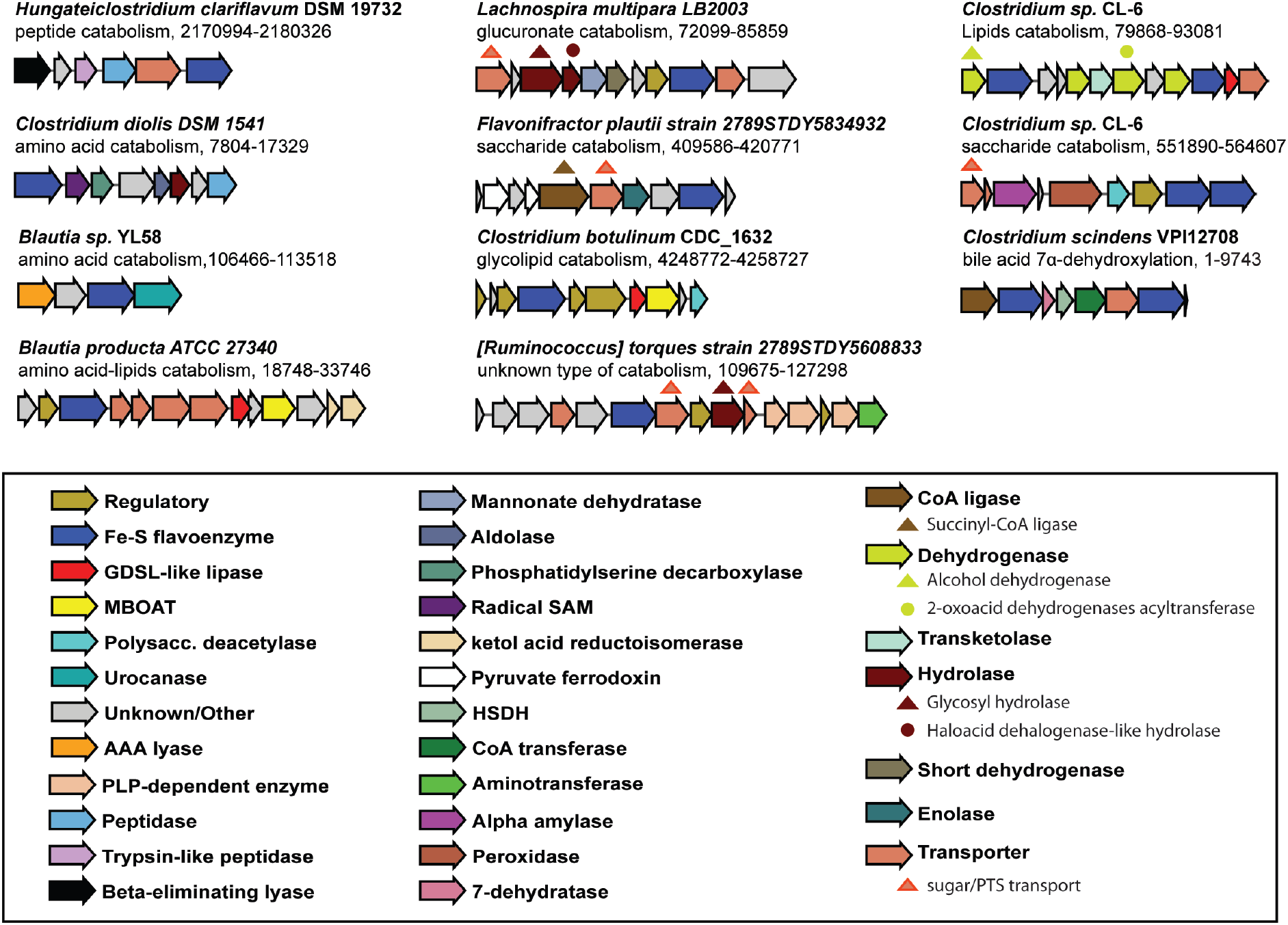
Diversity of Fe-S-flavoenzyme-encoding catabolic gene clusters from Clostridia. Gene clusters from Clostridia that contain a Fe-S flavoenzyme are numerous and diverse. Ten examples predicted to be involved in peptides/amino acids catabolism, saccharide catabolism, lipid catabolism and unknown type of metabolism have been picked from a collection of 1,052 GCFs, plus the *bai* operon. Genes are colored by predicted function.

## Conclusion

The metabolic potential of gut bacteria greatly exceeds the genetic potential of the human host. Therefore, bacteria can benefit from utilizing a diverse range of substrates that reach the digestive tract. Despite this, the mechanisms by which anaerobic bacteria catalyse these reactions are largely unknown. Here, we show that Fe-S flavoenzymes are likely to play a significant role in catalysing key redox steps within diverse catabolic pathways. Specifically, we found a remarkable expansion of Fe-S flavoenzyme copy numbers in the class Clostridia when compared to other gut-related bacteria. The strain-specificity of Fe-S-flavoenzyme-encoding gene cluster repertoires indicates that these enzymatic pathways may allow bacteria to specialize in the catabolism of specific dietary or host-derived molecules. We present a rich dataset of gene clusters that are candidates for detailed biochemical studies. Moreover, the presence or expression of these GCFs could be used as features alongside standard metabolic pathway annotations to assess whether their presence could explain variation in health/disease phenotypes, or whether they can explain variation observed in gut metabolomes^23,24^. All in all, given the importance of profiling the gut microbiome from a functional point of view, this study provides new ways of exploiting the genomic information present in public repositories, and provides a template for genomic exploration studies centred on key enzyme families to further understand the metabolic potential of the gut microbiome.

## Material and Methods

### Identification of members of the Fe-S flavoenzyme superfamily

BaiCD contains two Pfam domains, Oxidored_FMN (PF00724) and Pyr_redox_2 (PF07992). We used hmmsearch (HMMER 3.1b2, February 2015; http://hmmer.org/) to identify protein sequences harboring both domains from two databases: all bacterial sequences in GenBank (complete and draft genomes, >100,000 entries) and all sequences in RefSeq belonging to the class Clostridia (>2,500 entries), as some of the Clostridia genomes in GenBank lack gene coordinate annotations. Proteins with sequence e-value <= 1×10^−5^ for both domains were deemed hits. The resulting set of 49,437 Fe-S flavoenzyme superfamily members was used for the analyses described in the next three paragraphs.

### Construction of an Fe-S flavoenzyme protein similarity network

The amino acid sequences of the 49,870 Fe-S flavoenzyme superfamily members were clustered using MMseqs2^18^, setting the minimum identity to 0.9. From the 1,694 groups in the resulting protein similarity network, we picked five representatives of each cluster (randomly chosen when the cluster contained more than five nodes). ~2,500 singletons (families of size one) were also included in the subsequent analysis. We aligned the protein sequences to the Oxidored_FMN and Pyr_redox_2 Pfam domains using hmmalign (HMMER 3.1b2, February 2015; http://hmmer.org/), removed the unaligned and indel regions, merged the alignments of the two domains and reconstructed a phylogeny using FastTree^25^; the root of the tree was arbitrary placed. The resulting phylogenetic tree was annotated with interactive Tree of Life (iTOL^26^).

### Computational analysis of the prevalence and phylogenetic distribution of Fe-S flavoenzymes

We investigated the taxonomic distribution of Fe-S flavoenzyme genes and their variability in copy number; for simplicity, the number of hits per genome assembly was used as a metric of the copy number per genome. We constructed a phylogenetic tree of genomes in the class Clostridia using the following procedure: Clostridia genome assemblies harboring at least one Fe-S flavoenzyme gene were downloaded and quality-filtered using the N50 statistic, setting the threshold at 50 kb. The 16S rRNA sequences from the high-quality scaffolds were predicted using Barrnap version 0.9 (https://github.com/tseemann/barrnap). 16S rRNA sequences were aligned with Clustal Omega^27^, and FastTree was used to infer an approximately-maximum-likelihood phylogenetic tree. Finally, iTOL was used to display and annotate the phylogenetic tree. Two 16S sequences from *Bacillus subtilis* subsp. *subtilis* (NR_102783.2) and *Streptococcus agalactiae* DNF00839 (KU726685.1) were used as outgroups to root the tree.

### Analysis of the genomic context of Fe-S flavoenzymes

Using the subset of Fe-S flavoenzymes from the class Clostridia found in the RefSeq database, we inspected genomic context by identifying ‘neighboring genes’ that met the following criteria: they were encoded on the same strand as the Fe-S flavoenzyme and located within a maximum intergenic distance of 400 bp between subsequent genes. A script was used to parse the feature tables from the genome assemblies found to harbor at least one copy of the Fe-S flavoenzyme gene, from which the starting and ending coordinates of the cluster were extracted. Subsequently, the corresponding Genbank file of the flanking region was downloaded. The resulting Genbank file collection was used as an input for BiG-SCAPE^28^, which groups metabolic gene clusters (MGCs) into families. The output networks were visualized using the BiG-SCAPE interactive visualization tool.

## Data availability

Supplemental information for this article can be found in Zenodo repository (https://zenodo.org/) under this accession number: 10.5281/zenodo.3688515.

## Acknowledgments

This work was supported by the Chan-Zuckerberg Biohub (M.A.F.) and the U.S. Defense Advanced Research Projects Agency’s Living Foundries program award HR0011-15-C-0084 (M.A.F. and V.P.A.).

